# The threats to reptiles at global and regional scales

**DOI:** 10.1101/2023.09.08.556803

**Authors:** Harith Farooq, Mike Harfoot, Carsten Rahbek, Jonas Geldmann

## Abstract

At least 20% of reptiles are currently threatened with extinction. Due to the lack of comprehensive global assessments of the group, they have been omitted from spatial studies, especially the ones addressing conservation or spatial prioritization. With the recent release of global assessments for reptiles, in this study, we calculate the probability of seven biodiversity threats – logging, pollution, agriculture, invasive species, hunting, climate change and urbanization occurring at a global scale on a 50×50km grid and their impact measured as the probability of finding a threatened species. Despite the global nature of our approach, we compartmented the terrestrial land into 12 regions and conducted separate analyses on each region. We find that despite globally low levels of intensity, climate change and invasive species have the highest impact on reptiles. But when looked at the regional level, various threats have a high impact on biodiversity, even surpassing climate change and invasives. Our study highlights the importance of going beyond measuring the intensity of threats to measure impact and also the importance of subdividing the world into smaller units and testing the impacts separately.

## 1. Introduction

Reptiles are an important and often understudied taxa in nature conservation. They play a significant role in ecosystems (de Miranda, 2017) and can serve as indicators of environmental health, often responding more rapidly to human pressures than other vertebrate groups (Newbold, 2018). Understanding their needs and vulnerabilities is, thus, essential for designing effective conservation strategies to safeguard biodiversity. At least 1,829 out of 10,196 □ species (21.1%) of reptiles are threatened with extinction (IUCN, 2022) due to major threats including agriculture, logging, urban development, invasive species, and climate change (Cox et al., 2022). However, the lack of comprehensive assessments of reptiles by in the IUCN Red List of threatened species, has prevented the use of this group in global studies of biodiversity patterns and conservation priorities (Farooq et al., 2020; Farooq, Azevedo, et al., 2021; Fritz & Rahbek, 2012; Meyer et al., 2015; Rosauer et al., 2009). This is now changing. In the last years, we have seen major progress in the assessment of reptile species, in particular through the GARD initiative (Roll et al., 2017), through which the conservation status of over 10,000 species (∼ 99% of all described reptile species) have now been red listed. This comprehensive assessment of all reptile species has helped to highlight important differences in the spatial patterns of reptile richness and that of other vertebrate groups such as concentrations of rare species in unexpected locations such as parts of the Namib Desert or the Arabian Peninsula (Roll et al., 2017). As a consequence, many important areas for the conservation of reptiles may have been neglected in global conservation priorities (eg. Brooks et al., 2006; Strassburg et al., 2020).

Besides differences in richness patterns (Roll et al., 2017), reptiles may also differ considerably from the other groups of vertebrates in terms of threats, with several families being persecuted for fear of their venom (Gutiérrez et al., 2017) or for their meat and skin (Klemens & Thorbjarnarson, 1995), while other groups are especially attractive to the pet trade such as chameleons (Marshall et al., 2020).

Yet, simply understanding what species are under threat is not enough. To halt biodiversity loss and improve conservation responses we need to disentangle not only what are the threats to biodiversity but also where they occur. However, spatial information on threats to biodiversity remains limited and often only covering few of the many human threats affecting species (Joppa et al., 2016). This lack of data on threats remains one of the most important knowledge-gaps in conservation and one that significantly hampers our ability to cost-effectively respond to the current biodiversity crisis (Geldmann, 2023; Tulloch et al., 2015). To address this, we mapped the probability of impact for seven major threats to biodiversity: logging, agriculture, hunting, pollution, invasive species, climate change, and urbanization using the same approach as Harfoot et al. (2021). In addition, since the presence of a threat is not the same as species being significantly affected by those threats, we studied the relationship between the probability of a specific threat affecting a species with the probability of finding a threatened species. Our results provide the first global maps of threats to reptiles and provides important insights into how those threats can affect conservation priorities across the globe.

## 2. Methods

Range maps of reptile species were sourced from the IUCN Red List of threatened species (IUCN, 2022). In total, we used the ranges of 9,827 species, after removing all sea turtles (n = 6) and sea snakes (n = 48). For each species, we then identified threats listed as affecting individual species using the IUCN Threats Classification Scheme (Version 3.2). We focused on the five key drivers of biodiversity loss as identified by IPBES (2019): land use changes, direct exploitation of natural resources, climate change, pollution, and the invasion of alien species. Due to how the IUCN classifies threats; land use change was divided into agriculture, logging, and urbanization giving us a total of seven mapped threats (Table 1).

**Table 1.** Threat frequency and likelihood of impact.

| Threats | Number of species and percentage | Median LoI (SD) |
| --- | --- | --- |
| Logging | 1,313 (13.4%) | 0.04 (0.09) |
| Agriculture | 2,995 (30.5%) | 0.23 (0.19) |
| Hunting | 945 (9.6%) | 0.15 (0.15) |
| Pollution | 260 (2.6%) | 0.03 (0.07) |
| Invasives | 809 (8.2%) | 0.01 (0.08) |
| Climate change | 373 (3.8%) | 0.01 (0.05) |
| Urbanization | 1,387 (14.1%) | 0.09 (0.13) |
| Any | 4,551 (46.3%) | 0.38 (0.23) |

LoI = likelihood of impact.

We then used the species’ distribution ranges to build threat grid layers of 50 x 50 km for each of the seven threats. In these layers, each cell represents the probability of impact of a threat as the proportion of species occurring in that cell that is affected by a given threat while accounting for uncertainties introduced by the threat information being recorded at the species and not location level. This methods followed the approach used in Harfoot et al. (2021), using a binomial distribution where the response variable consisted of whether a species was coded as under a specific threat and weighted by the inverse of the cube root of range size. Also following Harfoot et al. (2021) we excluded any 50 x50 cells that had 10 or fewer species present in them.

For each 50 x 50 km grid cell, we also calculated the probability of finding a threatened species. We first classified all species as either ‘threatened’ (i.e. critically endangered (CR), endangered (EN), vulnerable (VU)) or ‘non-threatened’ (i.e. near threatened (NT), Least concern (LC)). We then applied the same methods as for assessing the probability of impact for individual threats, building on Harfoot et al. (2021), to estimate for each 50 x 50 km grid cell the probability of finding a threatened species. Since the larger the range map, the higher the uncertainty of the presence of a specific threat in a grid cell, we generated maps of uncertainty by calculating the median range size in each grid cell (Fig S4).

We then used linear models (LM) to test the relationship between the probability of impact from individual threats and the probability of finding a threatened species. For the relationship between the probability of impact from individual threats and the probability of finding a threatened species, we constructed models for both global and regional relationships. We used the IUCN regions (IUCN, 2022) for the regional analysis. This regionalization divides the world into 14 regions, of which we only excluded the Arctic and Antarctic from our analysis due to missing data (Table 2).

**Table 2.**
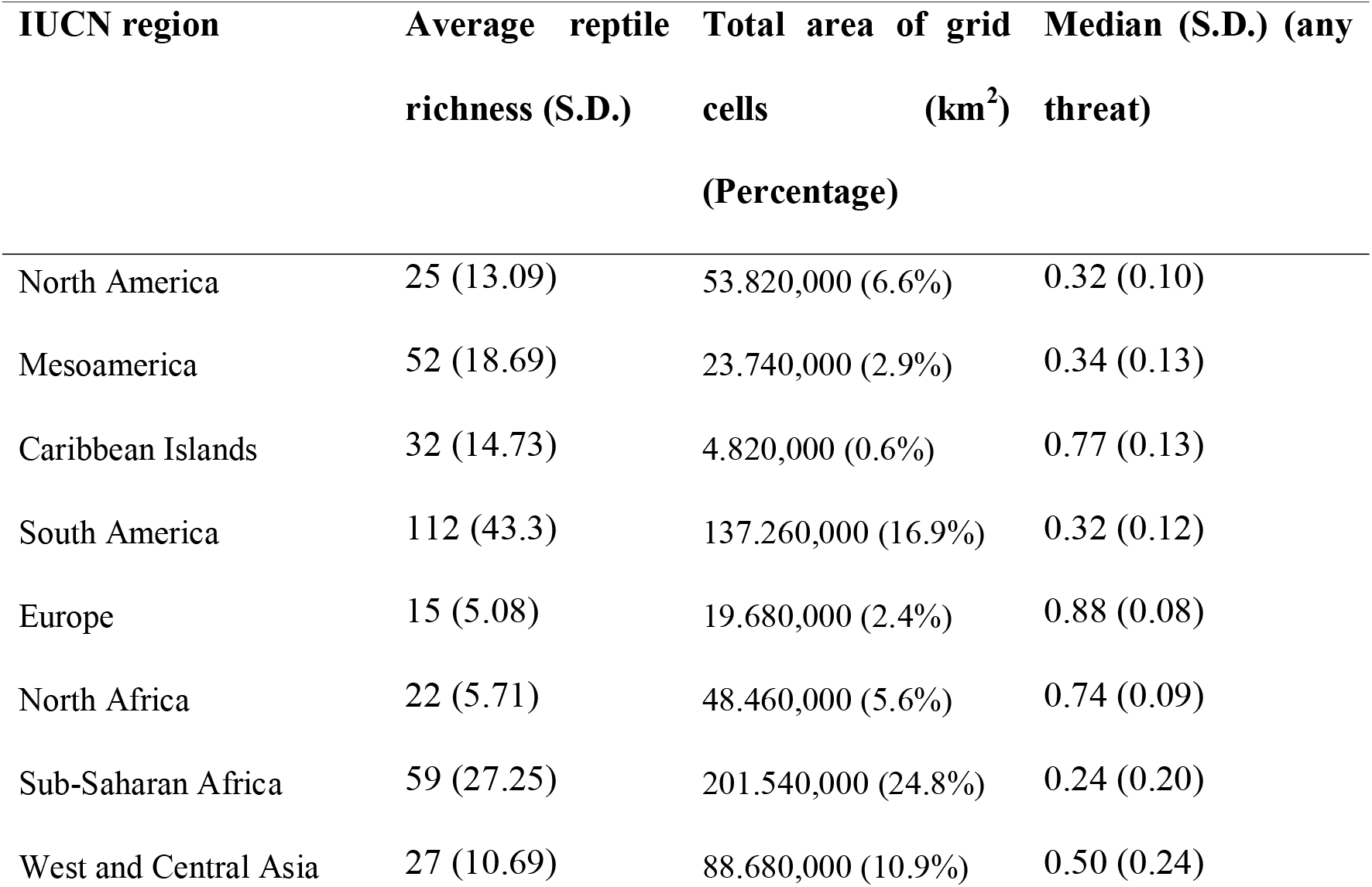
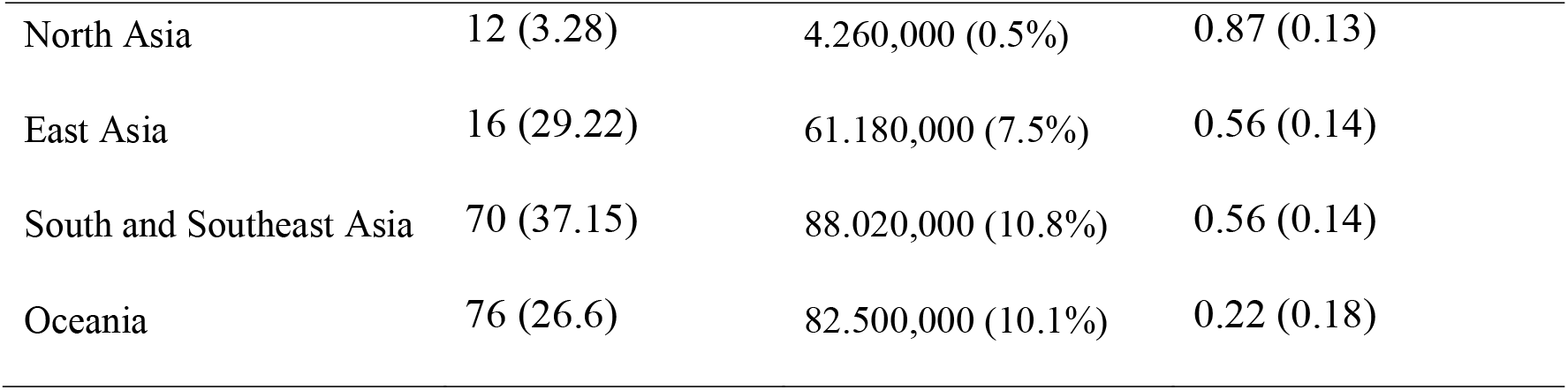
Geographic distribution of threats to reptiles. A table showing the number of average richness, total area of grid cells, and median threat probability with associated standard deviation for each of the analyzed IUCN regions. Values were calculated after removing all areas with less than 10 species. S.D. indicates the standard deviation of the estimates.

Finally, since the precision of our analysis is directly associated with the range size of each species, we generated maps of uncertainty by calculating the median range size in each grid cell (Fig S4). The higher the value, the higher the uncertainty of our results.

## 3. Results

Of the 9,827 species of terrestrial reptiles included in this analysis, 4,551 (46%), were affected by one or multiple of the seven threats. The threat that affected most reptile species was agriculture (n = 2,995; 31%), followed by urbanization (n = 1,387; 14%) and logging (n = 1,313; 13%; Table 1). However, the number of species affected by a particular threat did not always correspond to a comparably high median probability of impact. Thus, while hunting was only the fourth most reported threat to reptile species, it had the second highest median likelihood of impact, only superseded by agriculture and followed by urbanization, suggesting stronger spatial clustering (Table 1).

Across IUCN geopolitical regions, Europe (Median = 0. 88, Sd = 0.08) had the highest median probability of impact for any of the threats analyzed, followed by North Asia (Median = 0. 87, Sd = 0.13), the Caribbean Islands (Median = 0. 77, Sd = 0.13), North Africa (Median = 0. 74, Sd = 0.09), Southeast Asia (Median = 0. 56, Sd = 0.14) and East Asia (Median = 0. 56, Sd = 0.14; Table 2). Conversely, the region with the lowest median probability of impact of any threat was Oceania (Median = 0.22, S.D. = 0.18) and Sub-Saharan Africa (Median = 0.24, S.D. = 0,20; Table 2).

At the global scale, patterns for individual threat varied greatly (Figure 1). Thus, while agriculture was the most reported threat and that with the highest median impact, this was primarily driven by Europe, Central Asia, Madagascar, South America and the Caribbean islands (Figure 1c, 1h). Hunting dominated in Sub-Saharan Africa as well as in large parts of China and India (Figure 1e, 1h). Urbanization was primarily a concern in coastal regions and dominated coastal North America, Europe and North Africa, as well as the west coast of India and parts of the South American and Australian coastlines (Figure 1g, 1h). Higher probability of impacts from invasive species on reptiles was generally restricted to islands as well as Australia and New Zealand (Figure 1d, 1h). Logging was rarely the threat with the highest probability of impact, except for in New Guinea but often coincided with agriculture in other parts of the world (Figure 1a, 1c, 1h). Pollution (Figure 1b) and climate change (Figure 1f) showed low probability of impact for reptiles across most of the world.

**Figure 1.**
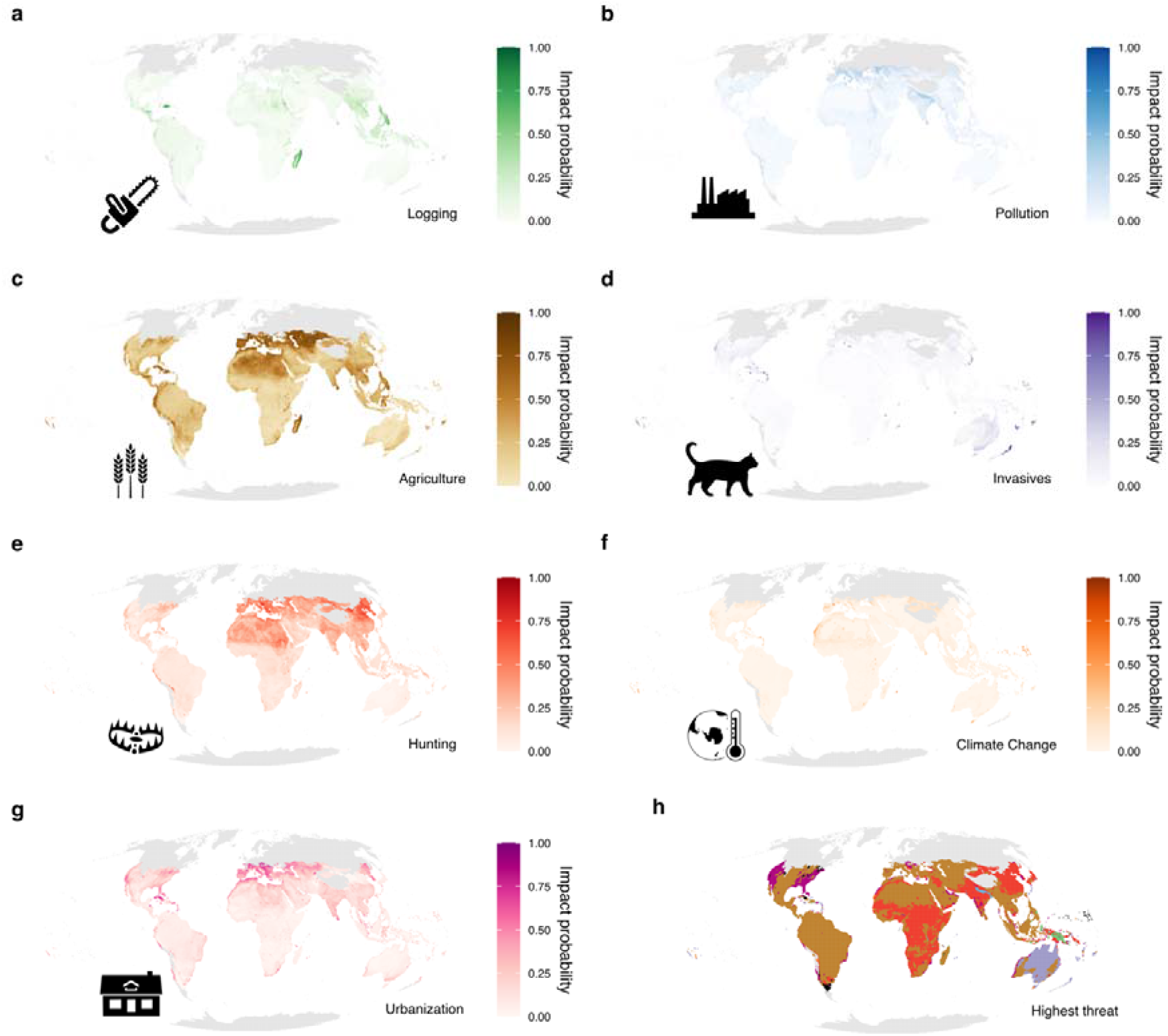
Global threats maps a – g, for logging, pollution, agriculture, invasive species, hunting, climate change and urbanization. In h, a map showing the highest probability of any of the threats in a – g, cells in black have an overlap of various threats as the ones with the highest probability. Grey indicates areas with 10 or less species for which a probability of impact was not estimated.

The relationship between the probability of impact of any threats and the probability of being a threatened species showed a positive and significant but low association when looking across all regions and for all threats (Figure 2). Looking at individual threats, the relationship between threats and a species being threatened was generally more similar within regions than within threats with the global south, and in particular South and Central America as well as Oceania experiencing the statistically most positive relationships between the probability of threats and the probability of being threatened (Figure 2). Conversely, for North America and Euro-Asia there were generally no or only weak relationships between threat and being threatened (Figure 2). Looking at individual threats, the relationship was stronger for invasive species, climate change and to a lesser extent agriculture indicating that for these threats the probability of threats was more associated with reptile species being threatened. The probability of being impacted by climate change had a more positive relationship with being a threatened species in North and Central America, Europe, Oceania and Sub-Saharan Africa while invasive species had a stronger association in South America, the Caribbean islands as well as in Oceania and Sub-Saharan Africa (Figure 2). Interestingly, the relationship between hunting and reptile species being threatened was negative for the Caribbean islands, indicating that areas with higher probability of hunting contained less threatened species. In the rest of the world, hunting was not strongly associated with threatened species, except for in South America, Oceania and Southeast Asia (Figure 2).

**Figure 2.**
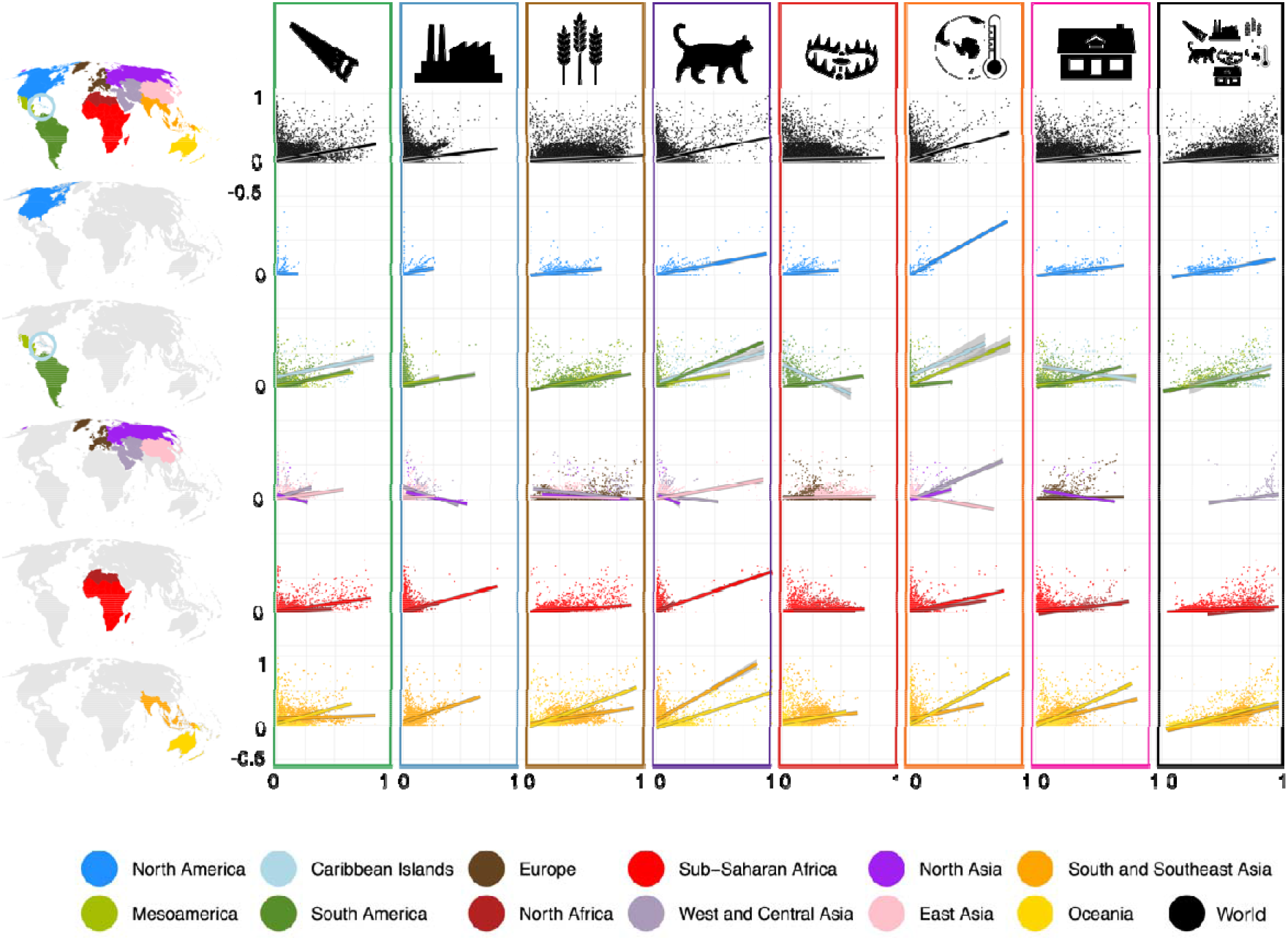
A figure showing the results of the linear model of the association between the probability of a species being threatened by the different threats (axis-x) and the probability of finding a threatened species (axis-y). Overall, there is a positive association between threatened species and all studied threats. When looking at the different threats and regions we see different threats affecting regions differently. We grouped the IUCN regions into biogeographic realms and connected the Australasian, Oceania and Indomalayan regions for visual purposes. Associations that were not significant were removed from the plot.

## 4. Discussion

### 4.1 Patterns of threat at the global scale

Using information from 9,827 species, we provide the first global maps of the probability of impact from the main threats to biodiversity for terrestrial reptiles (Fig 1). Our results show that while reptiles overall, are one of vertebrate groups under most pressure (Cox et al., 2022), this varies considerably geographically and across specific threats. Agriculture, urbanization, and hunting were the most pervasive threats (Figure S1). Agriculture and urbanization, often leading to habitat loss, are primary threats to reptiles as emphasized in non-spatial studies (Gibbons et al., 2000). Hunting poses significant risks, arising from the removal of reptiles from the wild for purposes such as food, “traditional” medicine, curios, and the pet trade. Additionally, reptiles face threats from unintentional by-catch during other harvesting activities, increasing instances of road mortality (Sparling et al., 2010), and persecution (Farooq, Bero, et al., 2021). These results also match expert opinion on reptiles which identified land use change, production and consumption, and human population as the main drivers for reptile loss (Isbell et al., 2023).

The Caribbean, Europe, North Asia, North Africa experienced the highest average probability of threat across all seven threats, followed by South and Southeast Asia and West and Central Asia (Figure S1). Hunting was a serious threat in South America, Southeast Asia, and Oceania. It is important to note that by hunting, we refer to IUCN’s threat code 5.1 – Hunting and trapping terrestrial animals. Therefore, not only the known consumption of reptiles ‘meat and eggs (Klemens & Thorbjarnarson, 1995) contribute to these high likelihoods in these regions, but also the collection of species for the pet trade (Marshall et al., 2020), the use of their skins (Wyatt et al., 2018) and also their persecution (Mesquita et al., 2015).

### 4.2 Linking probability of impact to threatened status

Our approach allowed us not just to detect where species are being impacted by human actions, but also to gauge how this impact potential relates to the risk of extinction. Recognizing that species can be affected by threats without necessarily being threatened, we integrated maps of impact likelihood with those showing the probability of locating a threatened species. This methodological increment of the analysis in Harfoot et al. (2021), enhanced our grasp of the interplay between threats and species conservation by highlighting focal areas and possible strategies to protect species under the risk of extinction. Overall, we found a weak but significant positive relationship between likelihood of impact and threatened species at the global level (Fig 2). Despite the unsurprising association between the exposure to threats and increased likelihood of extinction, the strength and even the direction of this relationship varied across IUCN regions as well as between threats. For instance, globally, the from the likelihood of being impacted by invasive species in a given pixel was strongly associated with the probability of being a threatened species (Fig 2). This was despite this threat activity exhibiting low median probability of impact (Fig S1). Thus, while invasive species might not affect a large number of species globally, where they do, they seem to be acutely linked to increased risks of extinction. The impact from invasive species is predominantly observed on islands in the Pacific, Caribbean, and Indian oceans. These ecosystems together with coastal mainland areas are hotspots of invasions (Pyšek et al., 2020). Invasive species are responsible for 14% of the critically endangered terrestrial vertebrates on land and 28% on islands, impacting at least 149 species and causing the extinction of ten (Dueñas et al., 2021). Adding to the high likelihood of threat of invasives in the Caribbean, we also found a strong positive association between the threat of invasive species and probability of being a threatened species in the region. This is likely linked to the invasive small Indian mongoose (*Urva auropunctata*) introduced between 1870 and 1872 to control the rat population (Pitt et al., 2017) but ended up responsible for the extinction of a number of species (Corke, 1992). They prey on a variety of items, including invertebrates, vertebrates, vegetable items or even human waste (Pitt et al., 2017). Included in its diet are reptiles and their eggs (Lewis et al., 2011).

Agriculture had the highest median probability of impact in any given grid cell across all regions (Fig S1). However, it also had a very low, and sometimes, even a negative, association with the probability of finding a threatened species (Fig 2). The only exception was Oceania, where we found a low average probability of impact for agriculture, but a strong positive association with the probability of finding a threatened species (Fig S1). These patterns underscore the critical necessity of comprehending the connection between threat probability and its repercussions on threatened species. This demonstrates that even though the likelihood of agriculture impacting a species’ population is low, the likelihood of encountering threatened species remains high. While determining causality is challenging, it remains undesirable to have any high level of threat probability within a grid cell where threatened species are found. Particularly since a single reptile might be susceptible to multiple threats (Gibbons et al., 2000).

We also found examples of negative relationships between the probability of impact and the probability of being a threatened species. These results highlight areas where despite the higher likelihood of impact from certain threats to reptiles, the areas have less threatened species. For example, in the Caribbean islands the probability of hunting affecting a species had a negative relationship with the probability of being a threatened species. This negative relationship is likely a result of the present distribution of the reptile species on the islands, where most species suitable for hunting have already been extirpated. This is confirmed by studies showing that up to 70% of reptiles have been lost in this region during the colonization of the island in the 1500s (Bochaton et al., 2021). Thus, given that our results are based on those species currently extant on the islands, they cannot capture past hunting pressure. Likewise, we find evidence for weak associations in overall threats in Europe despite high exposure to agriculture, pollution, hunting and urbanization. In Europe, this is likely a result of the present distribution of the reptiles in the continent, where species are being extirpated due to urbanization (French et al., 2018).

### 4.3 Limitations of the study

This study is based on layers of probability of threat that are based on IUCN Red List ranges coded for different threats (Harfoot et al., 2021). The advantage of these layers is that they consider the threat’s specificity to different taxonomic groups. When studying their association with the probability of finding a threatened species we generate a proxy for impact. Nevertheless, this approach suffers from three primary limitations. (1) The intrinsic uncertainty tied to IUCN’s range maps prevents us from performing the analysis at a high resolution (Di Marco et al., 2017). (2) It is also impossible for us to discern whether a particular threat directly leads to a species being classified as threatened, as it’s typically a culmination of multiple threats leading to the threat status. (3) Our study is also affected by potential uneven sampling (Hughes et al., 2021), underreporting (Cox et al., 2022; Winter et al., 2016) and lack of fieldwork to support the threat assessments in the IUCN Red List (Farooq, Azevedo, et al., 2021). In regions with good information, climate change, for example, is already the highest threat to reptiles, as seen in Europe (Falaschi et al., 2019) and the Caribbean Islands (Sinervo et al., 2010), a finding also supported by our analysis. In contrast, many other parts of the world characterized by high elevational heterogeneity, where reptiles are known to be affected by climate change (Moreira et al., 2023) are not showing up in our maps. Similarly, the Sahara Desert in Africa appears to be extensively affected by agriculture and hunting. This finding is, however, likely a consequence of the low richness of the region associated with the presence of large-ranged species (Fig S4c,e) such as the snake *Psammophis schokari* that expands across all of North Africa, through the Arabic peninsula and to Pakistan (IUCN, 2022) and is affected by agriculture in parts of the range other than the desert. The same holds true for the Desert Monitor (*Varanus griseus*) affected by hunting (IUCN, 2022). The high values observed in North Asia are likely inflated due to low species richness and small sample size (accounts for only 0.5% of the terrestrial cells used in the analysis). These effects of large range size on the uncertainty of our methodology can be seen in Figures S4a-h.

### 4.4 Conservation perspective

The understanding of which threats to biodiversity affect the various regions in the world is crucial to improving reptile conservation. Here we show that quantifying the probability of a specific human activity impacting a species provides valuable insights into the geography and diversity of threats affecting reptiles. However, as we also show here, such patterns do not always inform about how these impacts related to extinction risk. These patterns highlight the complicated relationship between the presence of threats and their impact. Thus, to improve our conservation responses we need to better understand both where threats are occurring and how they affect the biological communities in those areas.

The secretive behavior and characteristics of many reptiles, such as large home ranges, low population densities, and infrequent congregational tendencies, make it challenging to monitor their population trends. Consequently, populations might decline without immediate awareness. When a previously unmonitored species or population is identified as unexpectedly scarce or missing, the reasons behind their decline can remain elusive (Gibbons et al., 2000). To combat this, local universities and research centers should spearhead biodiversity documentation (Puruleia et al., 2023), addressing the gap in studies that comprehensively capture biodiversity threats. Intensive sampling campaigns with tailored methods are essential (Gonzalez et al., 2023). Such endeavors not only provide vital baseline data on national biodiversity but also bolster the scientific and conservation skills of budding professionals, while fostering community involvement as they are the enduring guardians of the land and its ecosystems.

## Supplementary materials

**Figure S1:**
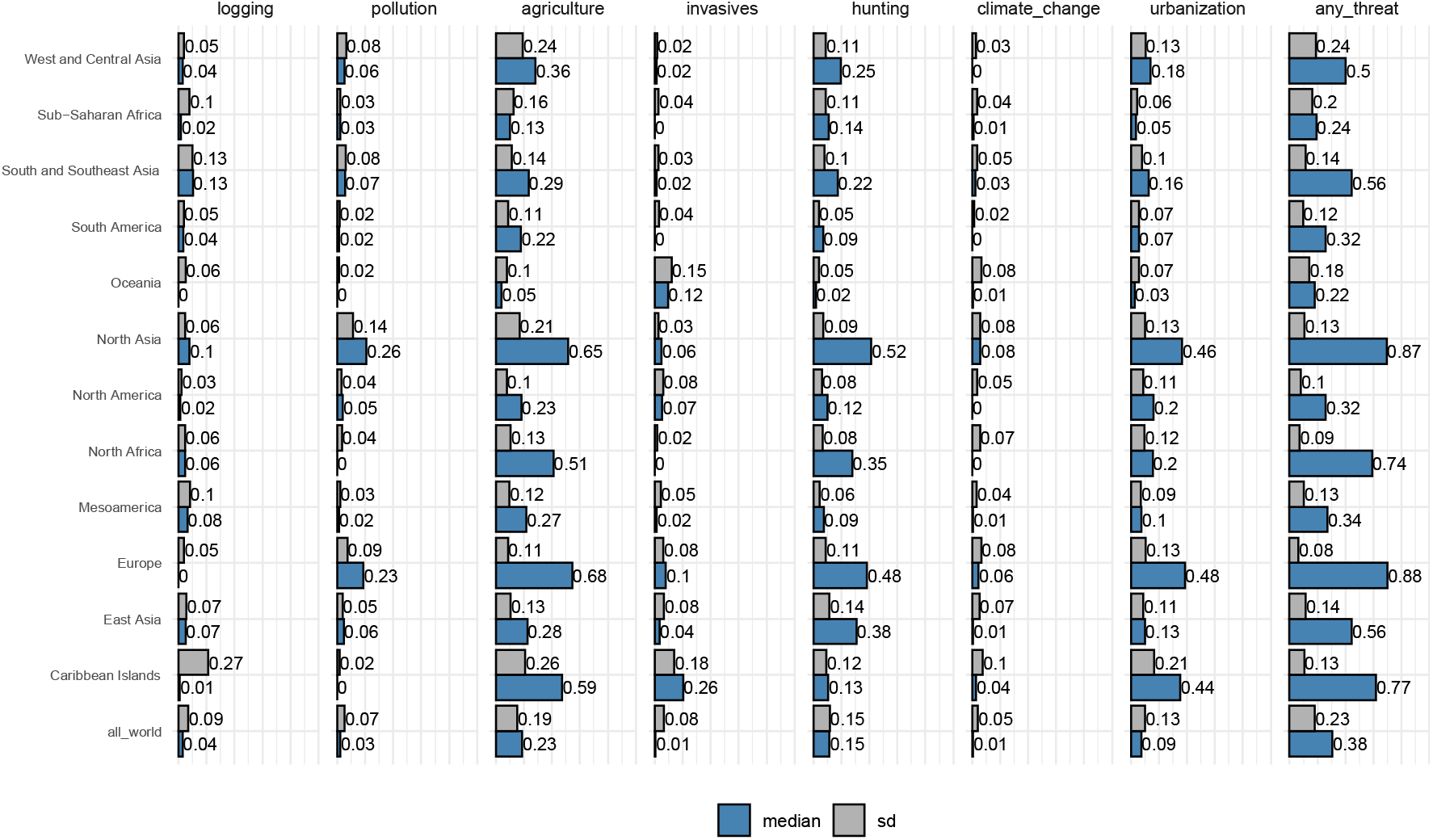
Median and standard deviation of the distribution of threat probability per region.

**Figure S2:**
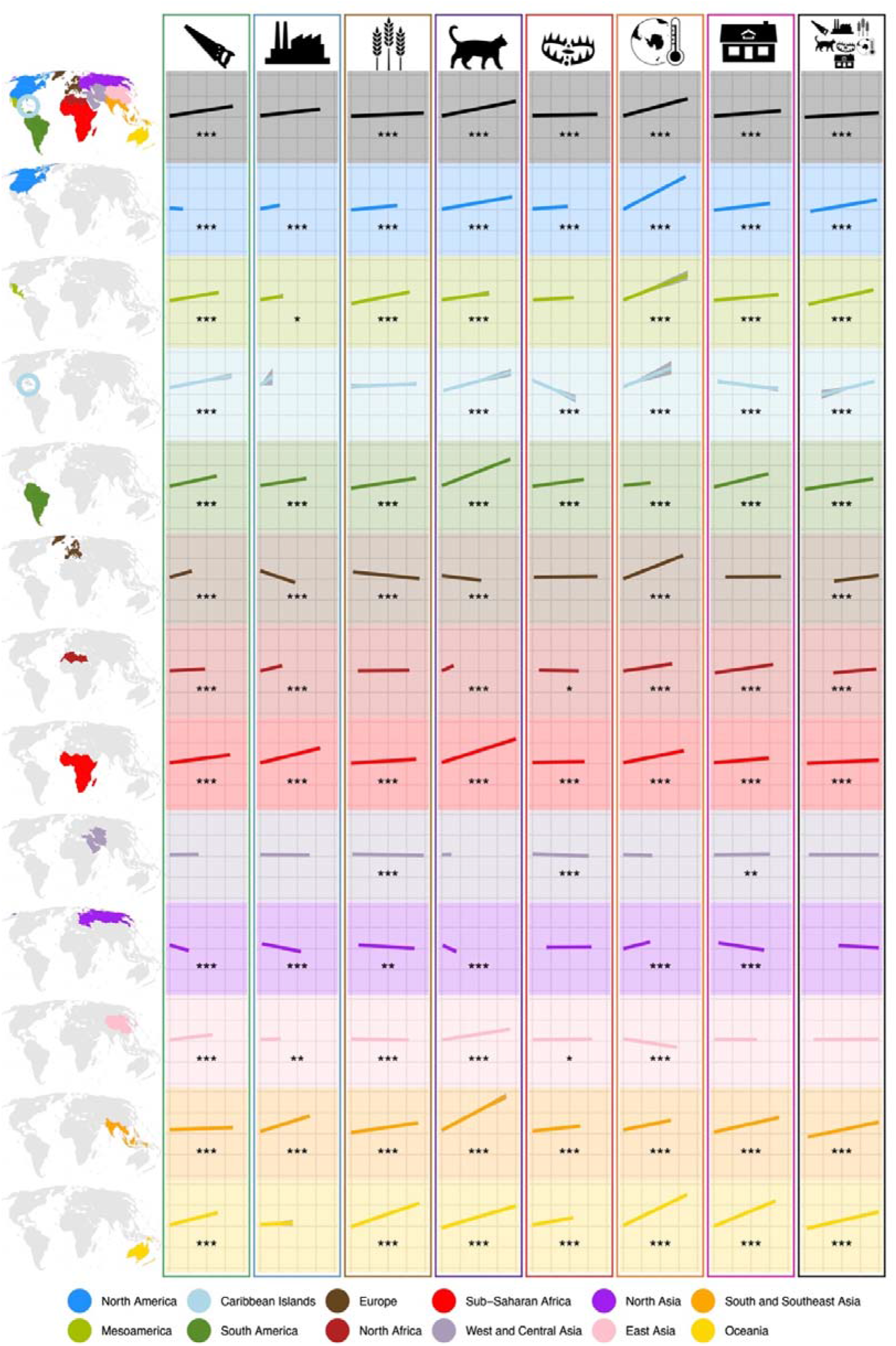
A figure showing the results of the linear model of the association between the probability of a species being threatened by the different threats. Overall, there is a positive association between threatened species and all studied threats. When looking at the different threats and regions we see different threats affecting regions differently.

**Figure S3:**
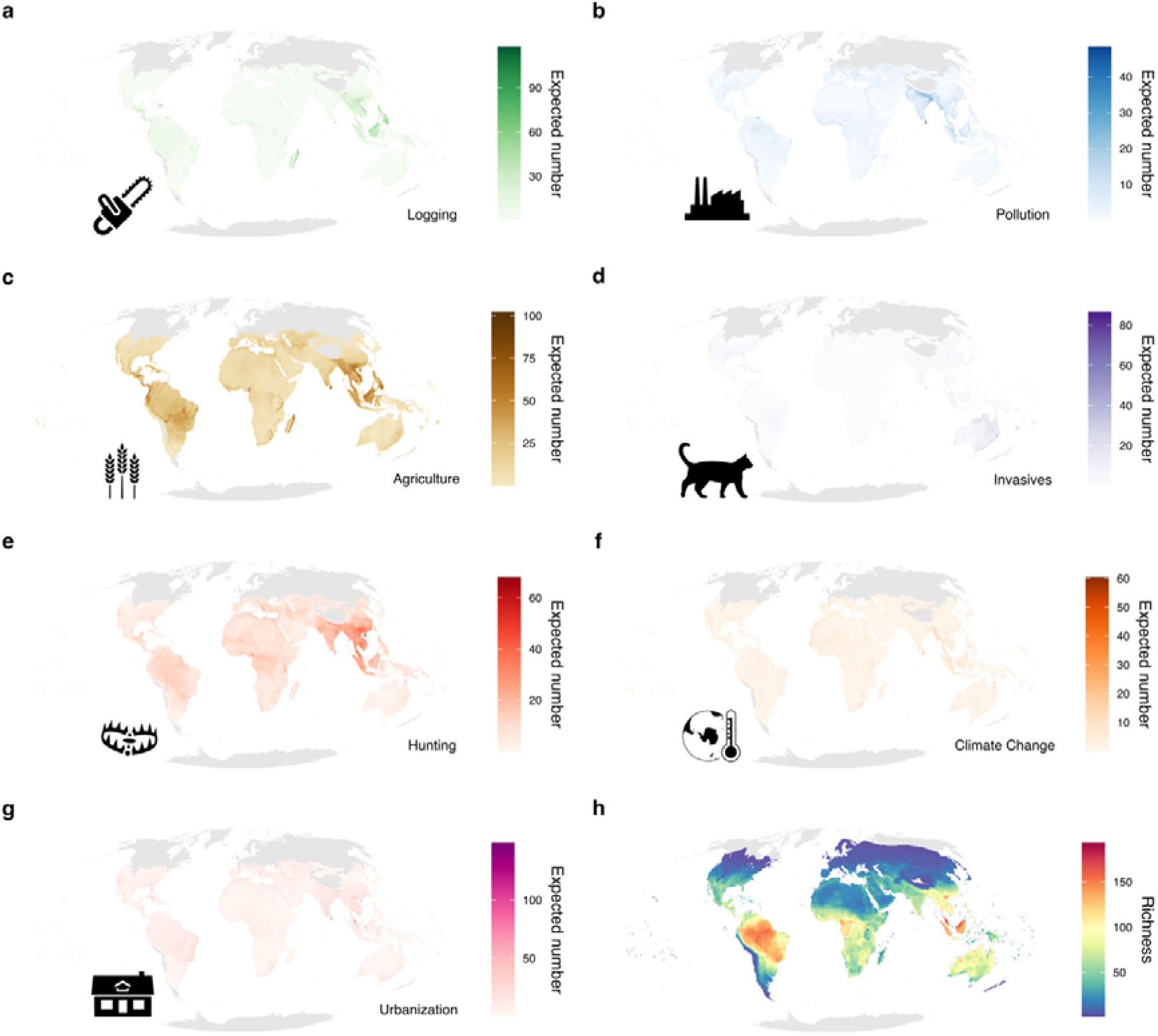
Global threats maps of the probability of impact multiplied by the richness in each grid cell: a – g, for logging, pollution, agriculture, invasive species, hunting, climate change and urbanization. In h the richness map for reptiles.

**Figure S4:**
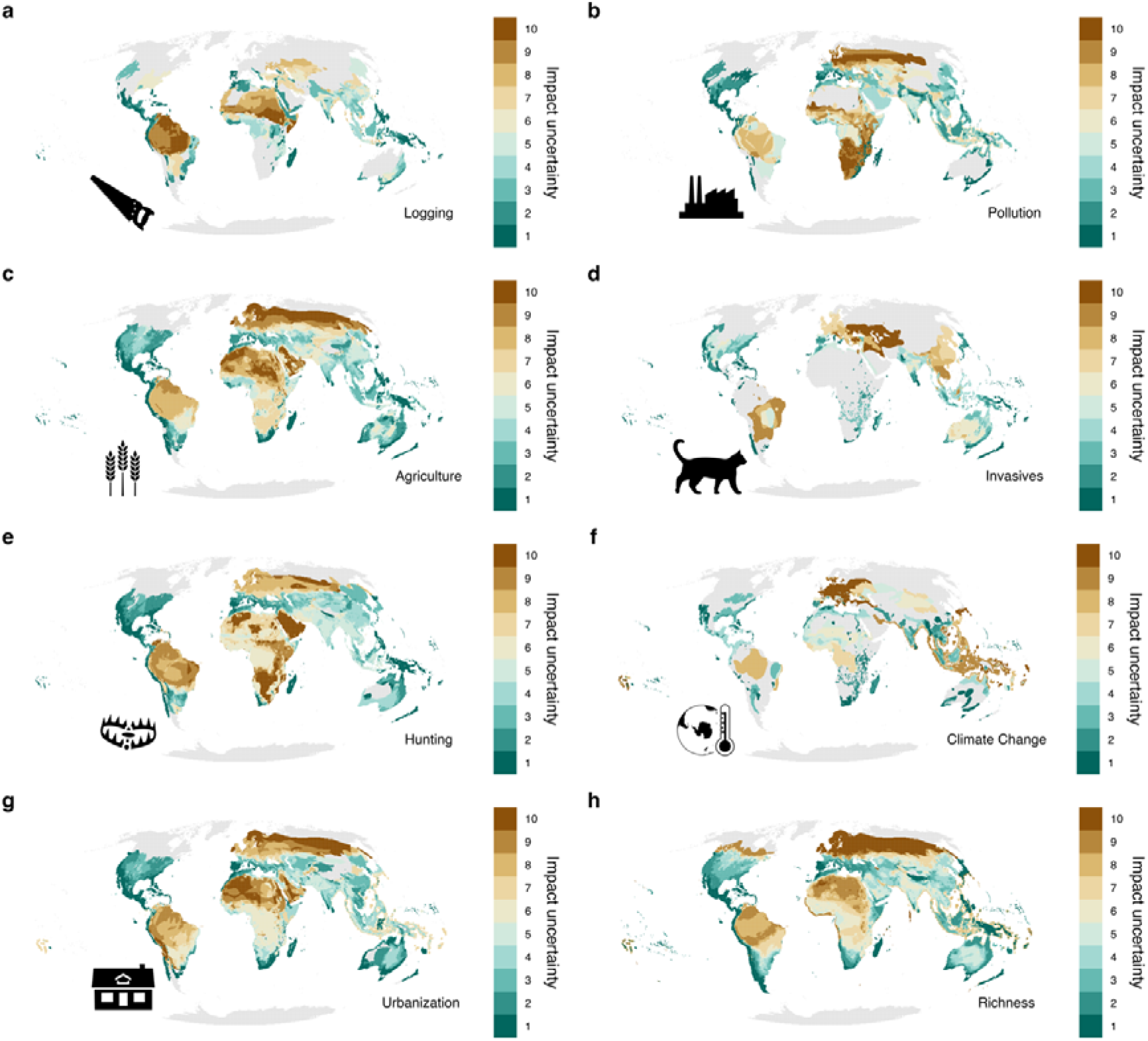
Global uncertainty maps of the probability of impact in each grid cell: a – g, for logging, pollution, agriculture, invasive species, hunting, climate change, urbanization and richness. For visualization purposes we log-tranformed the uncertainty and coloured the map by quantiles.

## Notes

### Competing Interest Statement

The authors have declared no competing interest.

